# Discrete synaptic states and context-modulated readouts support continual learning

**DOI:** 10.64898/2026.09.11.750704

**Authors:** Yuan Gao, Stefan Mihalas, Denis Turcu

## Abstract

Biological neural systems learn new behaviors while retaining earlier ones, whereas sequentially trained artificial networks often overwrite parameters and forget previously learned behaviors. We introduce the Context-modulated Synaptic-states Recurrent Neural Network (CoSyn-RNN), a new continual-learning model inspired by heterogeneous synaptic states and neuromodulatory control. CoSyn-RNN combines sparse recurrent allocation with a discrete protection rule: after each task, recurrent neurons whose synaptic weights cross a fixed threshold form a task-specific neuron group, and their associated parameters enter a protected state during later learning. A neuron-specific gain and a task-cued mask over a fixed base readout control how recurrent activity contributes to behavior. Across various task sequences consisting of many cognitive tasks, training and validation losses and accuracies were maintained throughout learning. The protection rule produced modular, task-ordered recurrent structure compatible with forward reuse of earlier computations. Pairwise recruitment measurements further defined minimum- and maximum-cost curricula that produced markedly different final recurrent structures and different remaining capacity for future learning. Thus, CoSyn-RNN provides a biologically inspired model in which protected synaptic states, sparse allocation, and context-dependent modulation support continual learning.

## 1 Introduction

Animals acquire new skills throughout life while retaining and reusing prior knowledge. This capacity is central to adaptive behavior, but its implementation is not well understood in biological circuits and remains unresolved in artificial networks. Several mechanisms, at different scales, could contribute. Synaptic tagging can separate transient from longer-lasting changes in synaptic efficacy^1^, neural activity associated with recent experience is re-expressed during subsequent sleep^2^, and contextual signals can restrict which pathways are active or plastic^3^. These processes are complementary: consolidation can protect selected changes, replay can revisit earlier activity, and gating can limit interference during new learning. Continual learning in a finite recurrent network raises two linked questions: can it retain earlier behaviors, and how does the order of experience affect the capacity left for new ones? The further challenge is to understand how a shared network allocates limited resources and whether earlier computations can contribute to later tasks^4^.

Biological work motivates complementary abstractions for protecting and expressing stored computations. Synaptic tagging distinguishes transient from longer-lasting changes in efficacy^1^, and models with multiple synaptic plasticity states can prolong memory^5^. Endoplasmic reticulum (ER) is distributed non-uniformly across dendritic spines and changes local calcium signaling and plasticity^6^. ER forms in active spines and can limit runaway potentiation, while larger spines can contain a spine apparatus formed from smooth ER^7^. The distinct calcium-dependent plasticity of ER-containing spines and the recruitment of ER to highly active spines make spine-apparatus synapses plausible candidates for a relatively stable or protected synaptic state^6,7^. This interpretation is a hypothesis, and it motivates asking whether heterogeneous plastic and consolidated states could help a circuit balance stability and flexibility. Additionally, at the system level, neuromodulation changes neuronal response properties and population dynamics^8^. In a computational model, a context-dependent signal can therefore serve as a functional analogue of a broadcast control signal that selects how shared activity is expressed.

Artificial neural networks expose the same stability–plasticity trade-off in a controlled setting. Sequential gradient-based training can overwrite parameters needed for earlier tasks, producing catastrophic forgetting (Fig. 1a). Existing approaches stabilize important parameters or previously learned functions^9–12^, allocate or prune task-specific capacity^13–15^, or constrain new gradients to reduce interference with previously learned solutions^16^. Context gating combined with synaptic stabilization already supports sequential learning of 20 cognitive tasks in a recurrent network^3^. Learned context-dependent gates can also recruit sparse neuronal ensembles in a feedforward model^17^. A thalamocortical model links contextual control to the protection of task-relevant circuits^18^. Other work has examined shared dynamical motifs and compositional reuse^16,19,20^. These studies establish retention and reuse as distinct goals. They leave room to ask how a model that recruits and protects recurrent neurons during learning distributes its remaining capacity across different task orders.

**Figure 1:**
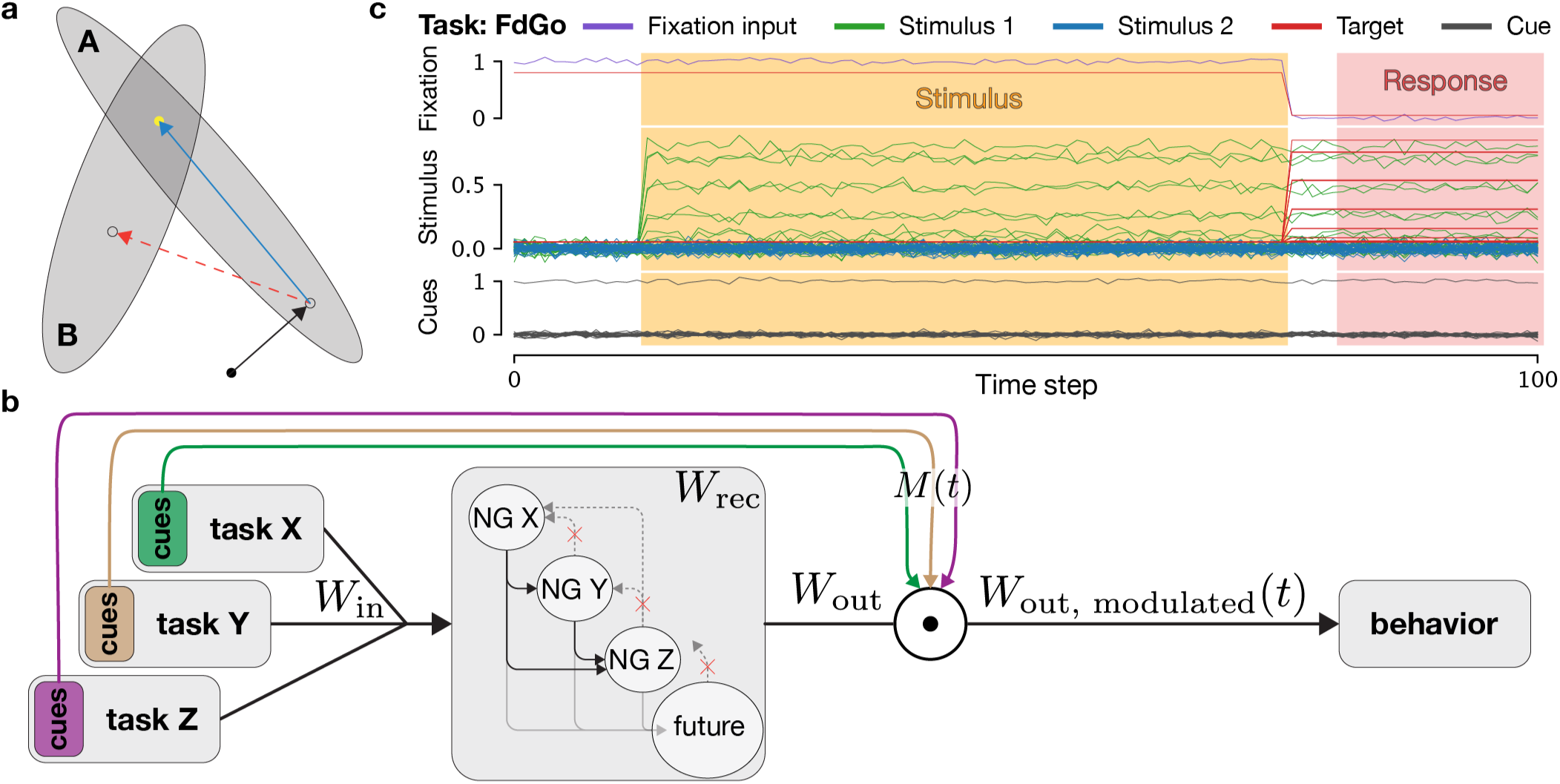
Schematic overview of CoSyn-RNN. **a** Conceptual schematic, adapted from^9^, illustrating two intersecting low-error parameter regions for tasks A and B. The black dot denotes the initial parameter state, and the yellow dot denotes a shared low-error solution. The black arrow illustrates learning task A from scratch. The red-dashed arrow illustrates how sequentially learning task B without protecting task A’s solution can move the parameters out of task A’s low-error region, resulting in catastrophic forgetting of task A. In contrast, the blue arrow illustrates how our model constrains subsequent parameter updates to remain within task A’s low-error region, thereby preserving performance on task A. **b** Schematic of the model architecture. Task inputs feed into the recurrent weights via the shared *Win* projection. The recurrent weights are optimized to preserve prior knowledge in task-dependent neuron groups (NG). The update rule permits connections from synaptically-consolidated groups into future learning, and connections into protected earlier groups are constrained and regularized, though not identically zero. Behavior is readout from the recurrent neurons via the task-cue-modulated projection *W*_out_ *⊙ M* (*t*). **c** Example trial from the `FdGo` task, illustrating the task input and target structure. Each trial contains a fixation channel, two population-encoded stimulus inputs, and a noisy one-hot cue signal. Full examples across tasks are shown in Supplementary Fig. S1.

Here, we introduce the Context-modulated Synaptic-states Recurrent Neural Network (CoSynRNN) as a framework for investigating how recurrent parameters can be allocated and protected during sequential learning (Fig. 1b). Sparse regularization and a neuron-specific gain promote compact recurrent solutions. After each task, a fixed weight threshold assigns recruited neurons and their incoming synapses to a protected state. Earlier groups may project into groups recruited later. A task-cued mask modulates a fixed base readout. We test sequential learning on cognitive tasks from the Yang et al. suite^21^ (Fig. 1c), measure the additional neurons recruited by each pairwise task transition, and use those directed measurements to select contrasting 16-task orders. CoSyn-RNN introduces a biologically inspired model for continual learning, where heterogeneous synaptic states maintain past learning while flexibly enabling future learning, and context-dependent modulation controls how recurrent computations are used efficiently. This design makes learning history and unused recurrent capacity observable aspects of the model. Its task-ordered connectivity permits forward influence from earlier groups, although the present analyses do not establish causal reuse of dynamics.

## 2 Methods

### 2.1 Model architecture

We used discrete-time recurrent neural networks to study sequential allocation within a shared recurrent population. The model combines leaky integration, sparse allocation, a neuron-specific gain, context-dependent readout modulation, and between-task synaptic state consolidation. Its dynamics are

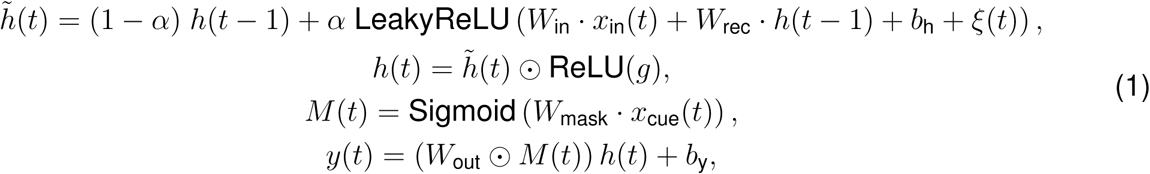

where *h̃*(*t*)*, h*(*t*)*, g ∈* ℝ*^H^* are pre-gain hidden activity, hidden activity, and gain for all hidden neurons *H*, *α* = *dt/τ* is the update rate and *τ* is the relaxation time constant, *x*_in_(*t*) *∈* ℝ*^F^* is the task input for all *F* input features, *x*_cue_(*t*) *∈* ℝ*^C^* is the task cue that encodes one of the *C* tasks as one-hot, *b*_h_ *∈* ℝ*^H^* is the hidden bias, *ξ*(*t*) *∈* ℝ*^H^* is 0-mean Gaussian noise with standard deviation *σ*_rec_= 0.05, *W*_in_ *∈* ℝ*^H×F^* is the input weight matrix, *W*_rec_ *∈* ℝ*^H×H^* is the recurrent weight matrix, *W*_mask_ *∈* ℝ*^O×H×C^* is the neuromodulation weight, *M* (*t*) *∈* ℝ*^O×H^* is the neuromodulatory mask applied to the non-trainable fixed readout weight matrix *W*_out_ *∈* ℝ*^O×H^*, *b*_y_ is the non-trainable fixed output bias, *y*(*t*) is the model output, “*⊙*” denotes element-wise multiplication, and “*·*” denotes dot-product. During training, independent 0-mean Gaussian noise with standard deviation *σ*_x_ = 0.01 is added to all input channels, i.e. *x*_in_(*t*) and *x*_cue_(*t*). We used a LeakyReLU nonlinearity to promote sparse activity without permanently silencing units after negative fluctuations early in training, and we constrained the neuromodulation mask between 0 and 1 using the Sigmoid function.

To coordinate recurrent allocation with the context-modulated readout during training, we included the nonnegative *g ∈* ℝ*^H^* state that scales the activity of each recurrent neuron. This acts as a learned neuron-specific gain without modifying the input or recurrent weight matrix. Biologically, neuromodulatory systems can alter neuronal gain and reorganize population dynamics, and low-dimensional neuromodulatory control in recurrent networks can support flexible context-dependent computation^8,22^. We use this correspondence only as a functional abstraction. *g* is not a mechanistic model of a particular transmitter, receptor, projection, or biological gain-control pathway. Although biological gain is constrained by intrinsic cellular properties, we did not need to impose an explicit upper bound on the gain parameter since the learned *g* values remained stable during training.

### 2.2 Task setup

We selected 16 tasks from the cognitive task suite of Yang et al.^21^. The task set includes Go, Anti, Decision Making (DM), and Delayed Decision Making (DelayDM) tasks. Each trial contains a fixation input, two population-encoded stimulus inputs carrying angular information, and a noisy one-hot cue that specifies the task context. The network generates the appropriate behavioral response based on the task cue. Figure 1c shows an example trial, and Supplementary Fig. S1 shows the input–output structure of all tasks. These tasks are useful for studying recurrent reuse because similar tasks can share dynamical motifs while retaining task-specific demands^19–21^.

### 2.3 Training and consolidation

We trained tasks sequentially using backpropagation through time and the Adam optimizer with learning rate 10^−4^, following related prior work^19–21^. The task objective was the mean-squared error (MSE) between the output and its target. To encourage each task to use a compact neuron group, we additionally regularized individual recurrent weights, neuron level incoming and outgoing recurrent connectivity, neuronal gain, and modulated readout. This allocation scheme is inspired by the group-sparsity regularization in dynamically expandable networks^15^. Additionally, since strong sparsity pressure may interfere with task acquisition before behavioral performance is established, the sparsity terms were scaled by the accuracy-dependent amplification factor

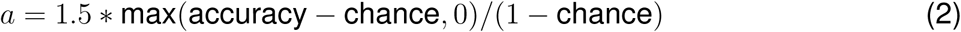

where chance and accuracy are defined in Supplementary Methods. The amplification increases as task accuracy increases, shifting optimization toward a sparser recurrent solution as the successful behavioral solution starts to emerge. We empirically set 1.5 as the maximum amplification through parameter sweeping to provide a trade-off between learning and sparsity.

For the vanilla control in Fig. 2, we trained a standard RNN without CoSyn-RNN’s synaptic consolidation or neuromodulatory readout. As in the recurrent model of Yang et al.^21^, inputs, including the task cue, entered through *W*_in_, and the output used a standard trainable projection rather than CoSyn-RNN’s task-modulated fixed readout. The training data and all applicable hyperparameters were the same as for CoSyn-RNN.

**Figure 2:**
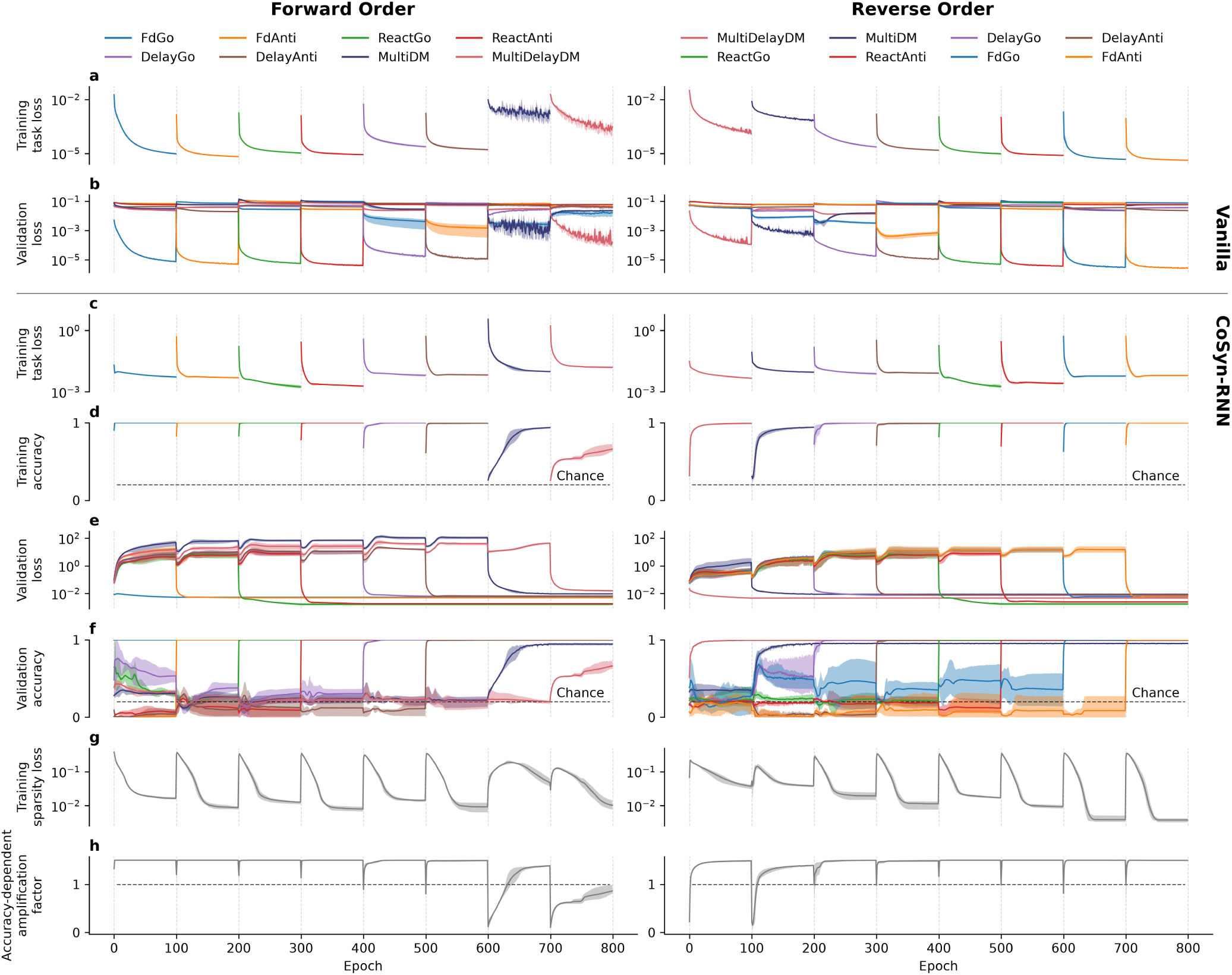
Continual-learning performance across task orders. The left column shows the forward order: `FdGo`, `FdAnti`, `ReactGo`, `ReactAnti`, `DelayGo`, `DelayAnti`, `MultiDM`, and `MultiDelayDM`. The right column shows the reverse order. For the vanilla RNN, **a** shows training task loss and **b** shows validation loss. For CoSyn-RNN, **c** shows training task loss, **d** training accuracy, **e** validation loss, **f** validation accuracy, **g** training sparsity loss, and **h** the accuracy-dependent amplification factor. Colors denote tasks, vertical lines indicate task transitions, solid lines show the mean across four random seeds, and shading shows the minimum-to-maximum range. Validation panels show all tasks throughout training, including tasks not yet introduced. Retention is assessed from each task’s curve after its training interval. Dashed lines indicate chance level in accuracy panels and unit line for reference in amplification panels. Training task loss is the MSE term in Eq. 3. Training sparsity loss is the sum of the remaining regularization terms. Validation shows task MSE without sparsity terms. Accuracy is defined in Supplementary Methods.

The complete objective was

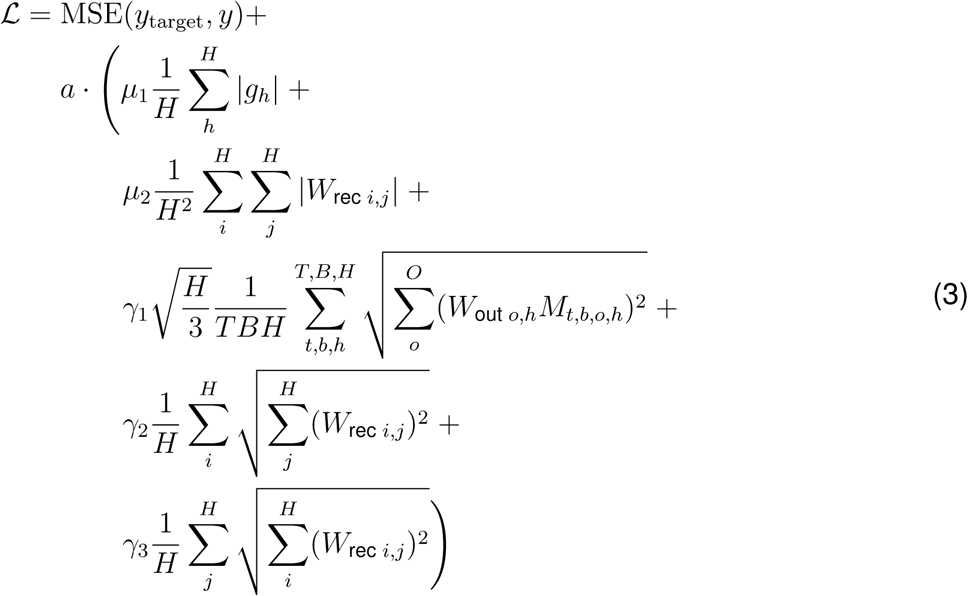

where *T ≈* 50-150, *B* = 256, *H* = 256, and *O* = 33 are the time, batch, hidden, and output dimensions, respectively. The time dimension was determined by the task definition and corresponded to approximately 1–3 s of simulated time. The batch size was chosen to provide stable optimization within the available computational resources, and the hidden dimension was set to *H* = 256, following prior work on the same or closely related tasks^19–21^. These prior studies used random task sampling and focused on recurrent networks with 256 hidden neurons, providing a network-size baseline for sufficient capacity. Here, *H* = 256 was also sufficient for sequentially learning all tasks while still leaving a subset of recurrent neurons available for future tasks. Thus, we did <u>not</u> vary the size of our models. The coefficients *µ_i_*and *γ_i_*weigh the regularizer terms, and *H/*3 compensates for the initialization scaling of the fixed readout weights. All reported experiments used *µ*_1_ = 0.1, *µ*_2_ = 1, *γ*_1_ = 0.1, *γ*_2_ = *γ*_3_ = 0.01, and a threshold of *θ* = 10^−4^. These values were selected through a parameter sweep based on validation performance.

After training each task, we consolidated the neuron group associated with that task. A trainable neuron was treated as inactive only if none of its incoming or outgoing recurrent weights and none of its context-modulated readout weights exceeded *θ* = 10^−4^ in absolute value. All active neurons for the current task were assigned to the learned task group. The recurrent synapses among neurons in the active group, together with all recurrent synapses incoming to that group, were then protected from updates during later tasks. Earlier formed groups could provide forward recurrent input to newly recruited groups. Our model imposes a discrete between-task consolidation step and is inspired by the distinction between transient and stabilized plasticity^1^, but it is not a mechanistic model of synaptic tagging, sleep, or replay. After consolidation and protection of just-learned-task synapses, all other recurrent synapses of the network became active for future tasks. During consolidation, we do not change or re-initialize any weights.

## 3 Results

### 3.1 CoSyn-RNN retains performance across sequential cognitive tasks

We first tested whether CoSyn-RNN could learn a sequence of tasks while preserving earlier solutions. We selected eight tasks spanning the task families described above and ranked them by empirical single-task difficulty, from easy (`FdGo`) to hard (`MultiDelayDM`). We call the easy-to-hard sequence the forward order and its reversal the reverse order. In the vanilla RNN control, training loss fell during each task’s own interval, but validation loss on previously learned tasks increased during subsequent training (Fig. 2a,b). This control illustrates forgetting in the tested standard RNNs that lack both synaptic consolidation and the task-modulated readout.

CoSyn-RNN learned each task as it was introduced in both orders (Fig. 2c,d). More importantly, the validation losses and accuracies for previously learned tasks remained stable as the model learned additional tasks (Fig. 2e,f). The learning trajectories differed between orders, but the final performance of most tasks changed little. Thus, the protected synaptic states and context-modulated readout maintained earlier behavior across sequential learning rather than merely fitting the current task.

Task order nevertheless affected how optimization balanced task performance against sparse allocation. The sparsity loss started high when a new task was introduced and decreased as the network adjusted to the new constraints (Fig. 2g). The dynamic balance between performance and sparsity, mediated by the accuracy-dependent amplification factor (Fig. 2h), shaped the learning dynamics and final performance of each task. We used an automatic amplification factor (Eq. 2), although its scaling may require tuning for tasks of different difficulty. The same tasks reached better final performance when learned early in the reverse order (compare ‘forward’ with ‘reverse’ in Fig. 2d,f for `MultiDelayDM`). This pattern suggests that task order influences how optimization balances task performance against sparsity, particularly for tasks of different difficulty, rather than exposing an inherent limitation of the continual-learning mechanism. Adjusting this balance according to task difficulty may preserve sparse allocation while ensuring that harder tasks retain access to sufficient recurrent resources.

### 3.2 Task-ordered recurrent structure and context-dependent readout

Ordering neurons by their number of above-threshold connections and marking their assigned group boundaries revealed a block-structured recurrent matrix (Fig. 3a), paired with corresponding active neurons (Fig. 3b) and the task-specific readout modulation (Fig. 3d–k) of the fixed readout base (Fig. 3c). Connections from later groups back to earlier protected groups were limited (above the block-diagonal), whereas the model permitted connections from earlier groups into later-recruited groups (below the block-diagonal). The sizes of assigned groups varied across tasks, and related Go and Anti family tasks often required smaller additions when one followed the other. While this does not directly prove forward reuse of dynamics, it suggests a structured allocation that could facilitate such reuse. Establishing reuse will require causal lesions or direct analysis of trajectories, subspaces, or fixed points^19^.

**Figure 3:**
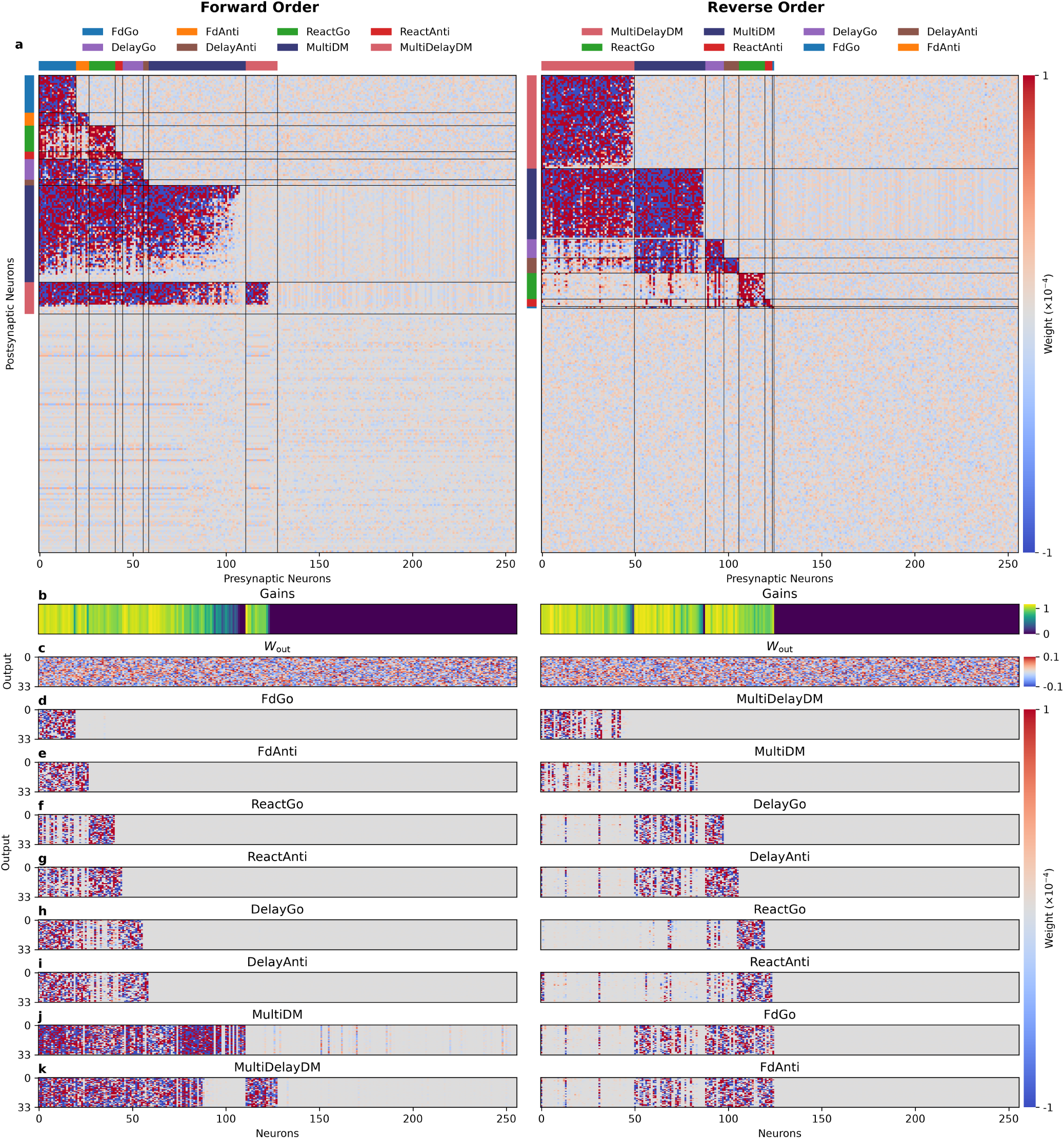
Recurrent structure and readout modulation after training. **a** Recurrent matrices *W*_rec_ for the forward- and reverse-order models. Neurons are sorted by the number of connections with 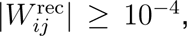 with a common order for both axes. Top and left colored strips and black lines indicate threshold-assigned groups and their boundaries. **b** Gains. **c** Fixed shared base readout matrices *W*_out_. **d–k** Task-specific context-modulated readout matrices. Weights are clipped to [*−*10^−4^, 10^−4^] for visualization purpose. Results for other random seeds are provided in the Supplementary Material.

The training curricula revealed a significant strength of the model’s sequential learning capabilities. CoSyn-RNNs have the ability to reuse previously learned dynamics for new tasks, reducing the need for additional task-specific recurrent neurons. For example, `FdAnti` recruited no additional recurrent neurons when it appeared last in the reverse order (Fig. 3a, right). Under this training procedure, an existing recurrent population and task-specific readout modulation were sufficient to express its solution. More broadly, the size of the neuron groups depended on the previously learned repertoire (Fig. 3a), generally increasing with task-group difficulty and decreasing in the reverse order. One interpretation is that earlier tasks established the available recurrent dynamics basis for later tasks, which are reused through the feedforward connections from previous blocks into these later groups. We explored this possibility further via task-order-dependent optimization in the next sections. Alternatively, the interaction between task loss and sparse regularization may also influence the observed allocation patterns, detailed below.

The two orders also revealed a trade-off between allocation and task performance. A substantial fraction of neurons remained unassigned after all eight tasks and thus available for future recruitment. In the forward order, `MultiDelayDM` acquired a small recurrent group (Fig. 3a, left) and reached slightly lower final accuracy than when it was learned first in the reverse order (Fig. 2f). This difference is consistent with an interaction between task difficulty and the loss– sparsity balance, but the present experiments do not isolate a causal effect of the regularizer. Further adjustment of that balance may improve performance without consuming excessive recurrent capacity, but exhausting that search space is outside the scope of this work.

### 3.3 Pairwise recruitment varies with task order

We quantified pairwise allocation for tasks in the Yang et al. suite^21^ by counting the additional recurrent neurons recruited when Task 2 was learned after Task 1 (Fig. 4a). This measure is directed because the recurrent state established by Task 1 determines the resources available to Task 2. For reference, the diagonal reports the number of neurons recruited when each task was learned independently. Longer sequences can change these pairwise estimates, but their number grows combinatorially. We therefore used pairwise recruitment as a first-order measure of task similarity.

**Figure 4:**
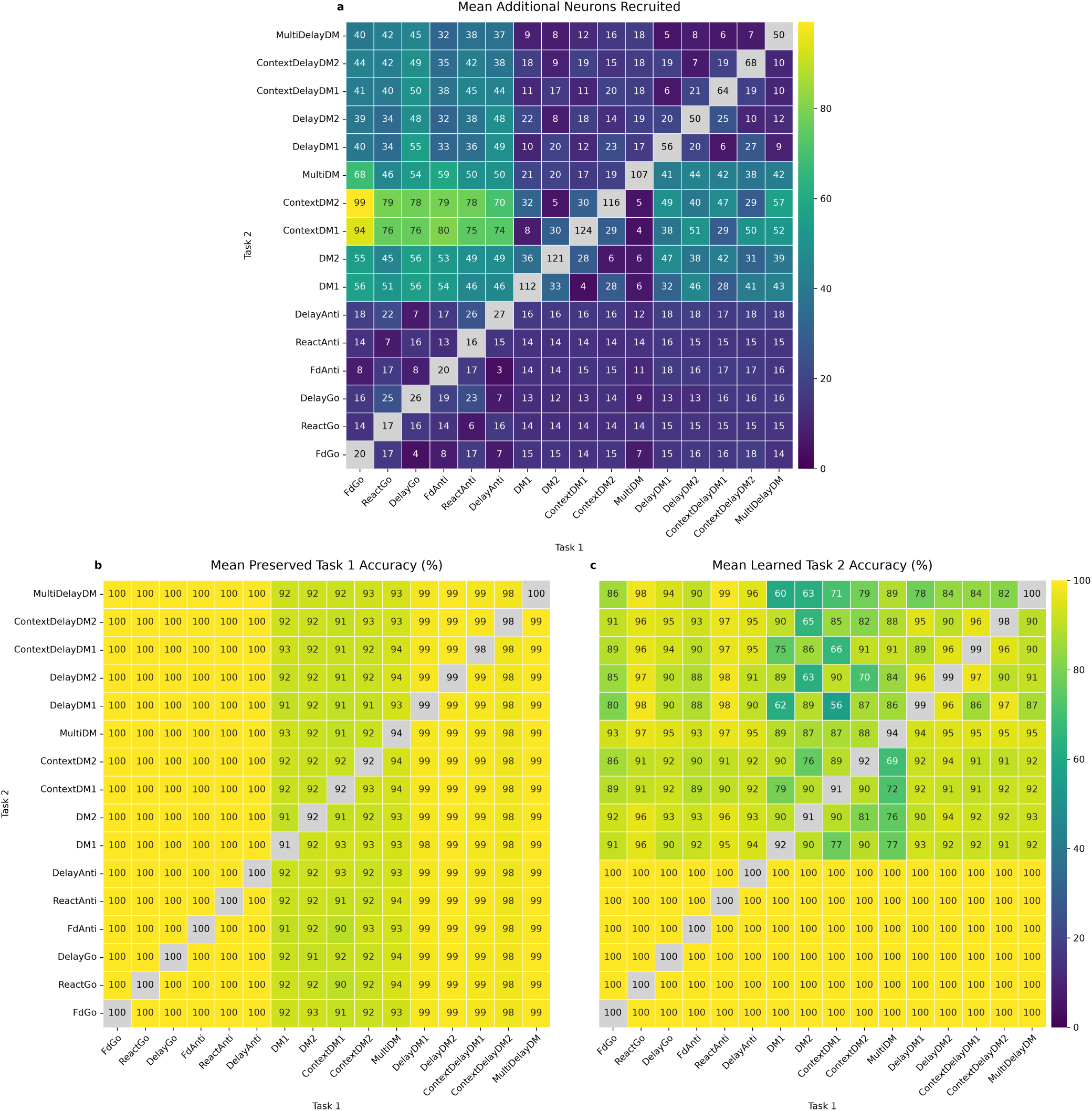
Pairwise recurrent allocation and task accuracy. Columns denote the first learned task (Task 1), rows denote the second (Task 2), and statistics use four random seeds. **a** Mean of additional neurons recruited for Task 2. Diagonal entries are the mean of the number of neurons recruited by the task independently. **b** Mean of preserved Task 1 accuracy after Task 2 training. Diagonal entries are the mean of accuracy learning task independently. **c** Mean of Task 2 accuracy after its training. Diagonal entries are the mean of accuracy learning task independently.

The pairwise matrix revealed structure across task families. Transitions from the Go/Anti family to the delay-DM family and from the delay-DM family to the DM family required moderately more additional neurons. By contrast, transitions from the Go/Anti family to the DM family required substantially greater expansion and formed the most prominent high-value region in the matrix. This pattern suggests that the dynamics established by Go/Anti tasks are less compatible with the computations required by DM tasks than with those required by delay-DM tasks. Recruitment differences may reflect the tasks’ computational demands and the effects of regularization, but do not establish incompatibility or reuse of particular recurrent dynamics.

By design, learning Task 2 did not disrupt Task 1 performance. Task 1 accuracy stayed nearly constant after learning Task 2 within each column, apart from variation across random seeds (Fig. 4b). Task 2 accuracy was also approximately constant within various row-blocks, indicating that it was governed largely by task complexity rather than by which task came first (Fig. 4c). Performance was particularly reduced when a DelayDM-family task preceded a DM-family task, suggesting less compatible recurrent dynamics between these task families in that direction. In contrast, performance was generally strong when a DM-family task preceded a DelayDM-family task, indicating a directional asymmetry in their interaction during sequential learning. This asymmetry may also reflect differences in how the two task families respond to sparsity pressure, since, as in Fig. 2, the learned solutions for the DM and delay-DM families were sensitive to the balance between task loss and sparsity regularization. This tradeoff warrants further investigation to determine how regularization can preserve sparse allocation without limiting task performance.

### 3.4 Curricula based on pairwise recurrent recruitment

The pairwise allocation matrix provides a tractable first-order proxy for the directional cost of learning one task after another, where cost denotes recruited neurons. Although training several tasks subsequently can alter these first-order estimates, they provide a useful guideline for constructing task orders that minimize or maximize the number of neurons involved in learning. Let *N_k_* denote the mean number of recurrent neurons recruited when task *k* is learned as the initial task, and let *A_kl_* denote the mean number of additional recurrent neurons recruited when task *l* is learned immediately after task *k*. For an ordered task sequence (*t*_1_*, t*_2_*, . . . , t_N_*) where each task appears exactly once, we defined the cumulative first-order allocation cost as

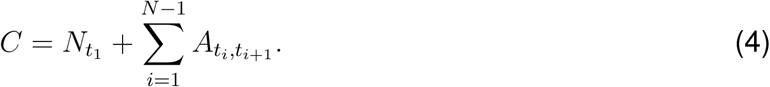

The first term accounts for the neurons required to learn the initial task, whereas the remaining terms approximate the additional neurons recruited during successive task transitions. We used a Held–Karp style dynamic programming algorithm^23^ to identify the minimum- and maximum-cost Hamiltonian paths (Supplementary Methods).

We trained separate CoSyn-RNNs on the complete sequence of 16 tasks using the minimum- and maximum-cost orders (Supplementary Methods). During each task’s training interval, its validation loss declined and accuracy rose significantly above chance. Performance then remained approximately stable as subsequent tasks were learned (Fig. 5a–f, similar to Fig. 2). The two curricula thus supported the full task repertoire but produced different final recurrent structures (Fig. 5g; Supplementary Figs. S6–S8). Consistent with the pairwise ordering proxy, the minimum-cost order tended to recruit fewer recurrent neurons overall, leaving a larger unassigned population available for subsequent tasks. Across four seeds, the final networks left 104 *±* 4 versus 62 *±* 5 neurons unassigned (out of *H* = 256). The contrasting structures provide the basis for different amounts of remaining capacity for future learning.

**Figure 5:**
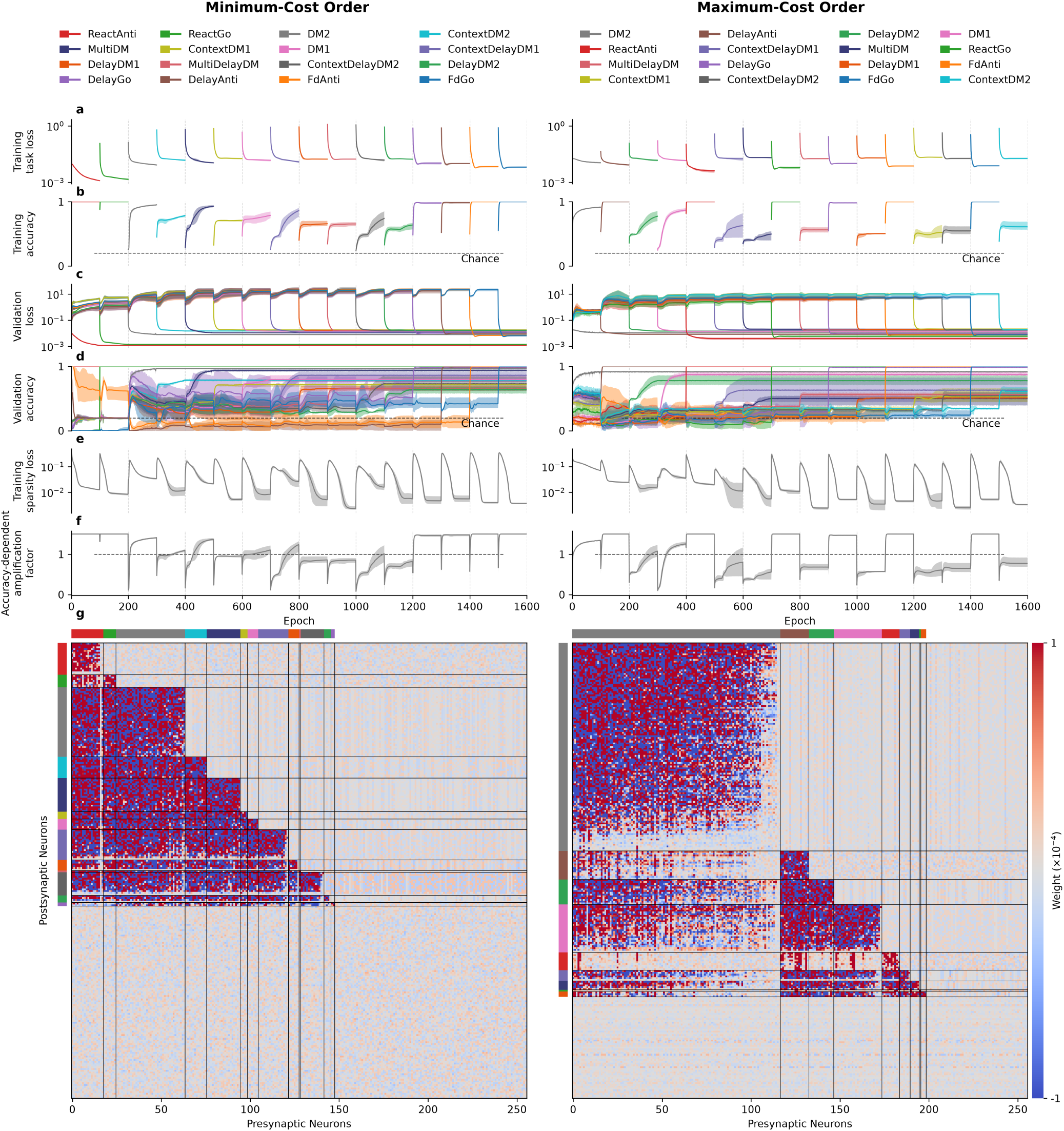
Performance and structure under minimum- and maximum-cost curricula. The left and right columns show the minimum- and maximum-cost orders, respectively. **a** Training task loss. **b** Training accuracy. **c** Validation loss. **d** Validation accuracy. **e** Training sparsity loss. **f** Accuracy-dependent amplification factor. Colors denote tasks, dotted vertical lines indicate task transitions, solid lines show the mean across four random seeds, and shading shows the minimum-to-maximum range. **g** Sorted recurrent matrices *W*_rec_ of the final models for seed `1`. Colored bars indicate task-associated groups and black lines mark group boundaries. Corresponding results for other seeds appear in Supplementary Figs. S6–S8. Validation panels include tasks before their training interval. Retained performance should be read from each task’s curve after training.

The minimum-cost order did not follow an intuitive progression based on task family, empirical difficulty, or complexity. Instead, it tended to place difficult tasks relatively earlier than most simpler tasks (see order of all 10 variants from DM-family in Fig. 5b, left), consistent with the previous eight-task comparison in which harder tasks were easier to learn before sparsity and protection constrained the network. This pattern is qualitatively consistent with previously proposed optimal ordering principle where tasks should be learned from the least representative to the most typical^24^. However, this ordering differs from the biological intuition behind behavioral shaping, where simpler behaviors often scaffold more complex ones, but it does not contradict that principle. Rather, it points to different learning mechanisms underlying artificial and biological learning systems. The model uses explicit task boundaries, gradient-based optimization, and a fixed balance between task loss and sparsity, all of which differ from biological learning.

The observed structures also depend on hyperparameters, particularly the balance between task loss and sparsity regularization. Exhaustively searching that space is not computationally feasible and is beyond the scope of this study, and different settings could alter both performance and recurrent recruitment. The selected curricula should therefore be interpreted as a first-order demonstration that task order can change how much recurrent capacity remains for future learning, not as a universal difficult-first learning rule.

## 4 Discussion

CoSyn-RNN combines plastic and protected synaptic states with context-modulated output control in a recurrent continual learning model. In the tested cognitive-task sequences, it retained earlier solutions as new tasks were learned, whereas the vanilla RNN control forgot them. The resulting connectivity reflected the intended task-ordered constraints, and the current results suggest causal computational reuse. Recruitment of neurons between pairs of tasks revealed task-specific allocation patterns that depended on the order of task presentation. Such allocation allowed us to infer a first-order approximation of how to best train our recurrent model while maintaining flexibility to learn future tasks.

This contribution is biologically inspired and builds on earlier recurrent continual-learning models. Previously, context-dependent gating combined with synaptic stabilization preserved performance across cognitive tasks^3^, learned gating also recruited task-related ensembles in a feedforward model^17^, a thalamocortical model used contextual control to protect task-relevant circuitry^18^, and other models have examined how recurrent computations can be organized or composed across tasks^16,20^. CoSyn-RNN differs in its learned, threshold-assigned neuron groups, its task-modulated fixed readout, and its use of directed recruitment to compare task orders. We deliberately do not compare performance between the prior models and our approach, focusing instead on novelty of computational mechanisms and biological motivation. Synaptic tagging, ER-dependent plasticity, and replay motivate distinct timescales and states of plasticity^1,2,6,7^, and CoSyn-RNN assigns threshold-defined neuron groups to a protected set at explicit task boundaries. Likewise, cue-dependent readout masking captures a computational function that neuromodulation could support^8,22^, although we do not model a specific transmitter, receptor, projection, or multi-area circuit.

Several limitations define the reach of these results. The synaptic-state threshold treats participation as binary and may consolidate weak but dispensable weights or exclude weak but important ones. Sparse regularization schedule interacts with task difficulty and could compress a new solution prematurely. Broader robustness remains to be tested across alternative task suites, network sizes, or hyperparameter ranges. Finally, additional neuron recruitment given learned tasks is a measure supporting dynamical motif reuse, but the mechanisms of this process require further insights. Analyses of hidden-state geometry, trajectory alignment, fixed points, task-dependent dimensionality, and causal lesions are needed to determine what computations are shared.

Future work could replace the fixed synaptic-state threshold with an adaptive importance estimate, allow graded or reversible consolidation, and compress several consolidated groups without replaying the original training data. The last direction would require a principled way to sample or preserve the network’s internal activity rather than assuming access to past examples. Additional analyses can test whether earlier groups causally support later tasks through lesions, trajectory comparisons, and fixed-point or subspace analyses. Higher-order analyses could also quantify how an entire learned repertoire, rather than the immediately preceding task, changes the cost of acquiring the next task. Extending the model to interacting recurrent areas with explicit neuromodulatory pathways would directly test the cross-area interpretation suggested by the functional readout gate.

Together, our results show that continual learning in a recurrent network can be supported by coordinating three functions: protecting previously recruited synapses, allocating unused recurrent capacity to new tasks, and using context to control how shared dynamics drive behavior. CoSyn-RNN learned a diverse suite of cognitive tasks sequentially while preserving earlier performance, and its task-ordered connectivity and directed recruitment costs revealed how learning history shapes both network structure and the capacity remaining for future tasks. By assigning concrete computational roles to heterogeneous synaptic states and context-dependent modulation, CoSyn-RNN provides a tractable framework for connecting biological hypotheses about stability and plasticity with artificial continual-learning methods. More broadly, this work suggests that durable learning need not require either complete parameter stability or unconstrained plasticity, but can emerge by protecting established computations while selectively recruiting and expressing new ones.

## 5 Acknowledgements

We thank Haozhe Shan and the Mihalas group for helpful discussions and feedback. YG and DT were supported by the Shanahan Family Foundation Fellowship at the Interface of Data and Neuroscience at the Allen Institute and the University of Washington, supported in part by the Allen Institute. SM was in part supported by NSF (Nos. 2223725 and 2424124) and NIH (Nos. R01EB029813 and RF1DA055669) grants. This research was supported by the Allen Institute, founded by Jody Allen – chair and co-founder of Allen Family Philanthropies, and the late Paul G. Allen – investor, philanthropist, and co-founder of Microsoft. We gratefully acknowledge their vision and generosity, which make this work possible.

## 6 Supplementary Material

### 6.1 Code availability

All training and analysis code is publicly available at https://github.com/ivanygao/CoSyn-RNN.

### 6.2 LLM use statement

ChatGPT/Codex was used as an assistive tool during code development, figure preparation, and manuscript editing. For programming, it helped generate repetitive code and identify implementation errors. It also assisted with visualization scripts and figure aesthetics. The model architecture, training pipeline, analysis framework, and associated methodological decisions were designed and developed by the authors. For manuscript preparation, it was used for grammar checking and language polishing and rephrasing. The manuscript content, including the scientific flow of the ideas and results presented, the relevant literature background, and the key discussion points, was designed and written by the authors. All scientific decisions, analyses, interpretation of results, and final manuscript content were reviewed and determined by the authors.

### 6.3 Supplementary Methods

#### 6.3.1 Task Sets

The 16 tasks used in this paper were adapted from the cognitive task suite of Yang et al.^21^. Here, we summarize the task-specific fixation, stimulus, and response requirements used in our experiments. The full input–output structure is shown in Supplementary Fig. S1.

##### Go and Anti task families

The Go- and Anti-family tasks each presented a single stimulus in either modality 1 or modality 2. Go tasks required a response in the direction of the stimulus, whereas Anti tasks required a response in the opposite direction. The three variants differed in the temporal relationship between stimulus presentation, fixation offset, and response initiation.

`FdGo, FdAnti`. A single stimulus was presented in either modality 1 or modality 2 before the fixation cue was removed. The network responded in the stimulus direction for FdGo and in the opposite direction for FdAnti.
`ReactGo, ReactAnti`. A single stimulus was presented in either modality 1 or modality 2 while the fixation input remained active throughout the trial. The network was required to respond immediately after stimulus onset, either in the stimulus direction for ReactGo or in the opposite direction for ReactAnti.
`DelayGo, DelayAnti`. A brief stimulus was presented in either modality 1 or modality 2 and was followed by a delay period that ended when the fixation cue was removed. The network then responded in the remembered stimulus direction for DelayGo or in the opposite direction for DelayAnti.

##### Decision Making (DM) and Delayed Decision Making (DelayDM) task families

In all DM- and DelayDM-family tasks, stimulus 1 was sampled uniformly between 0*^◦^* and 360*^◦^*, and stimulus 2 was sampled between 90*^◦^* and 270*^◦^* away from stimulus 1. The network was required to report the direction of the stronger stimulus according to the task-specific modality rule. In the DM tasks, the two stimuli were presented simultaneously, whereas in the DelayDM tasks, they were separated in time.

`DM1, DM2.` Two stimuli were presented simultaneously and remained available until the end of the trial. Both stimuli appeared in a single sensory modality. They appeared in modality 1 for DM1 and in modality 2 for DM2. The network reported the direction of the stimulus with the larger amplitude.
`ContextDM1, ContextDM2`. Two stimuli were presented simultaneously in both sensory modalities and remained available until the end of the trial. For ContextDM1, information from modality 2 was ignored and the network reported the direction of the stronger stimulus in modality 1. For ContextDM2, information from modality 1 was ignored and the network reported the direction of the stronger stimulus in modality 2.
`MultiDM`. Two stimuli were presented simultaneously in both sensory modalities and remained available until the end of the trial. The network integrated stimulus strengths across the two modalities and reported the direction of the stimulus with the larger combined strength.
`DelayDM1, DelayDM2`. Two stimuli were presented at separate times in a single sensory modality. They appeared in modality 1 for DelayDM1 and in modality 2 for DelayDM2. The network compared their amplitudes and reported the direction of the stronger stimulus.
`ContextDelayDM1, ContextDelayDM2`. Two stimuli were presented at separate times in both sensory modalities. For ContextDelayDM1, information from modality 2 was ignored and the network reported the direction of the stronger stimulus in modality 1. For ContextDelayDM2, information from modality 1 was ignored and the network reported the direction of the stronger stimulus in modality 2.
`MultiDelayDM`. Two stimuli were presented at separate times in both sensory modalities. The network integrated stimulus strengths across the two modalities and reported the direction of the stimulus with the larger combined strength.

#### 6.3.2 Accuracy

We evaluated response accuracy on a held-out validation set after each training epoch. All tasks used the same evaluation rule. For each trial, the population-coded ring output was decoded and evaluated at the final valid time point of the response phase, defined as the last time point at which the output mask was nonzero. A trial was classified as correct when the minimum circular distance between the decoded and target response angles was less than 36*^◦^*, following the criterion used by Yang et al.^21^. Under a uniformly distributed response angle, this tolerance corresponds to a chance accuracy of 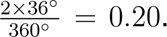 Each task had 128,000 training trials, and evaluation was performed using 4,096 held-out trials per task.

#### 6.3.3 Training

All models used a time step of *dt* = 20ms, a relaxation time constant of *τ* = 100ms, and *H* = 256 hidden units. Recurrent weights were initialized from a uniform distribution and rescaled to have an initial spectral radius of 1. Additive 0-mean Gaussian recurrent noise with standard deviation *σ*_rec_ = 0.05 was applied during training. Recurrent activity used a LeakyReLU activation function with negative slope 0.01. The neuron-specific gain used a ReLU activation, the modulation used the sigmoid activation with gain 20, and the output activation was linear.

Models were trained using Adam with a learning rate of 10^−4^ and mean-squared error as the task loss. Default Adam parameters were used. The total training objective included a gain sparsity coefficient *µ*_1_ = 0.1, a recurrent sparsity coefficient *µ*_2_ = 1, a modulated-readout coefficient *γ*_1_ = 0.1, and recurrent incoming- and outgoing-connection coefficients *γ*_2_ = *γ*_3_ = 0.01. The synaptic-state threshold was set to *θ* = 10^−4^. Each task’s 128,000 training trials were split into 500 mini-batches of 256. Every example was used once per epoch, and training continued for 100 epochs per task. Hyperparameters were selected through a parameter sweep.

#### 6.3.4 Task ordering

We used a Held–Karp-style dynamic-programming algorithm^23^ to identify the minimum-cost and maximum-cost Hamiltonian paths under the objective in Eq. 4. For each subset of tasks *S* and terminal task *j ∈ S*, the algorithm stores the optimal cumulative cost of a path that visits every task in *S* exactly once and ends at *j*. A single-task path ending at task *j* is initialized with cost *N_j_*, representing the number of recurrent neurons recruited when *j* is learned first. For larger subsets, the optimal cost of ending at *j* is obtained by considering each possible predecessor *i ∈ S \ {j}*, taking the stored optimal cost for a path through *S \ {j}* ending at *i*, and adding the pairwise transition cost *A_ij_*. The minimum-cost solution selects the smallest value at each recurrence, whereas the maximum-cost solution selects the largest. After all tasks are included, backtracking through the stored predecessors recovers the corresponding task order. Because the objective includes the initial-task allocation and only adjacent pairwise transition costs, while omitting dependence on the complete preceding task history, the resulting orders are first-order approximations of the full sequential-learning process.

Using the task indices

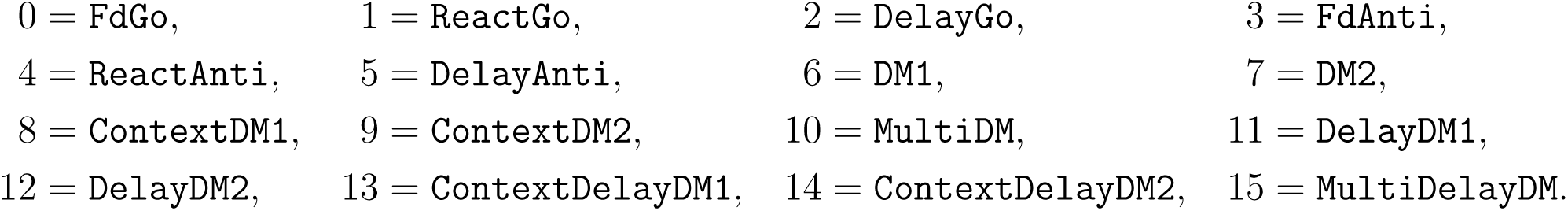

|  |  |  |  |
| --- | --- | --- | --- |
| 0 = FdGo, | 1 = ReactGo, | 2 = DelayGo, | 3 = FdAnti, |
| 4 = ReactAnti, | 5 = DelayAnti, | 6 = DM1, | 7 = DM2, |
| 8 = ContextDM1, | 9 = ContextDM2, | 10 = MultiDM, | 11 = DelayDM1, |
| 12 = DelayDM2, | 13 = ContextDelayDM1, | 14 = ContextDelayDM2, | 15 = MultiDelayDM. |

the minimum-cost order was

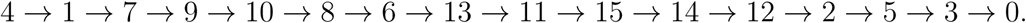

with allocation cost 174.25. The maximum-cost order was

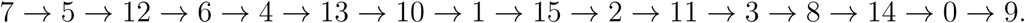

with allocation cost 695.0. Note this cost is larger than our model size of 256 recurrent neurons. Nevertheless, when training all tasks sequentially, our model uses less than 200 neurons out of the total networks size of 256, further supporting reuse of dynamics from previously learned tasks.

### 5.4 Supplementary Figures

**Figure S1:**
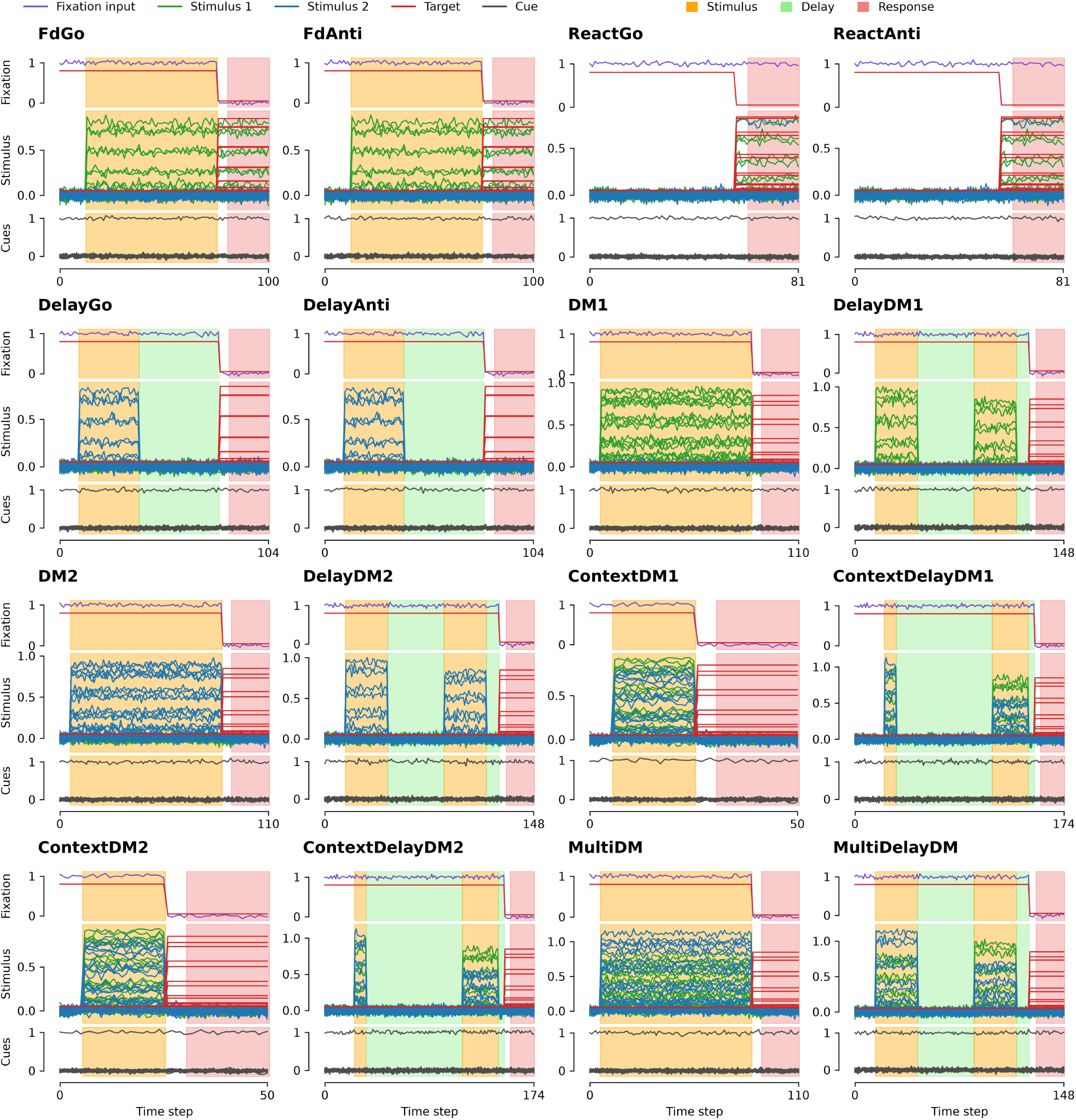
Example trials for all 16 tasks.

**Figure S2:**
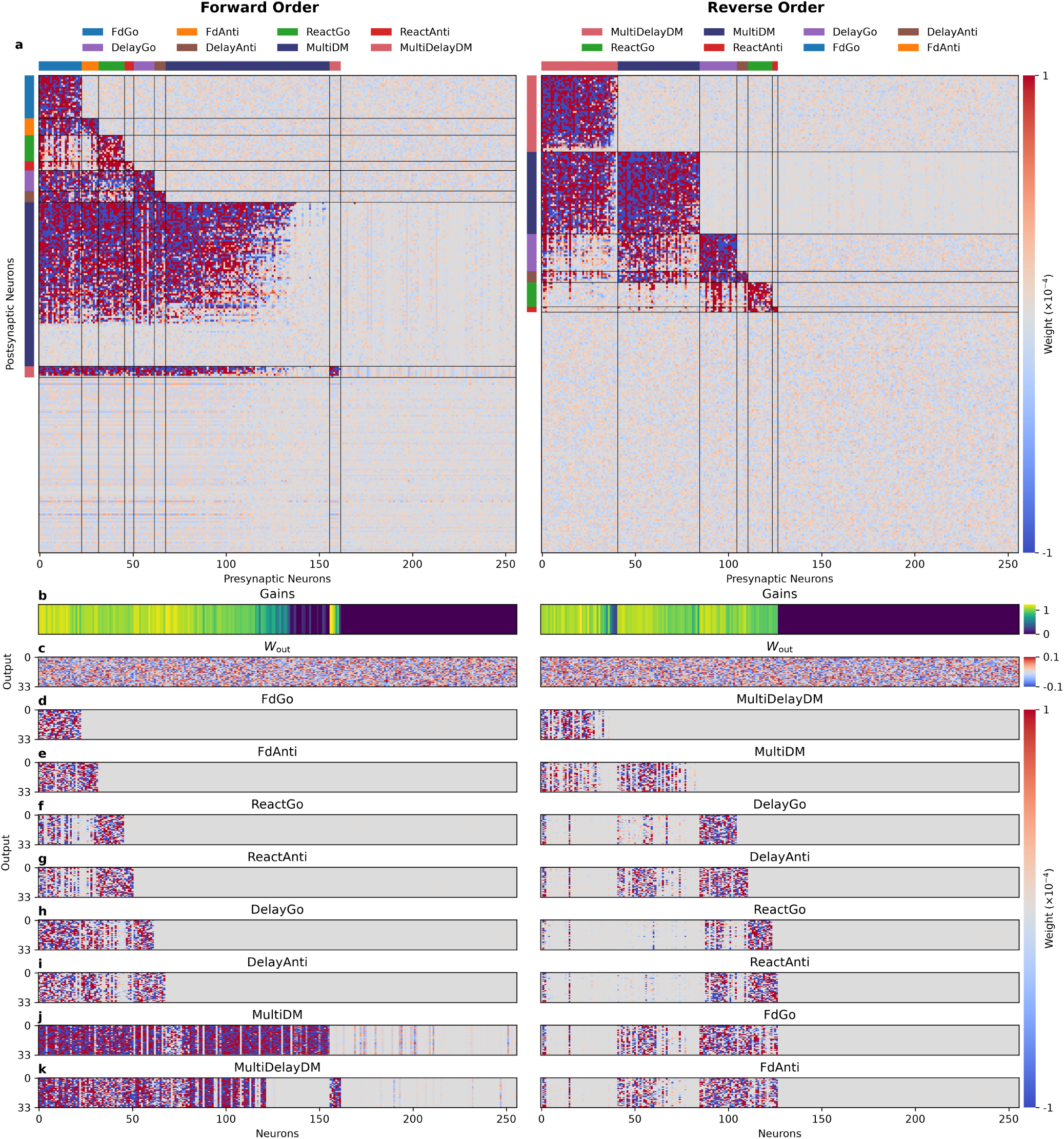
Same as Fig. 3, for random seed 42.

**Figure S3:**
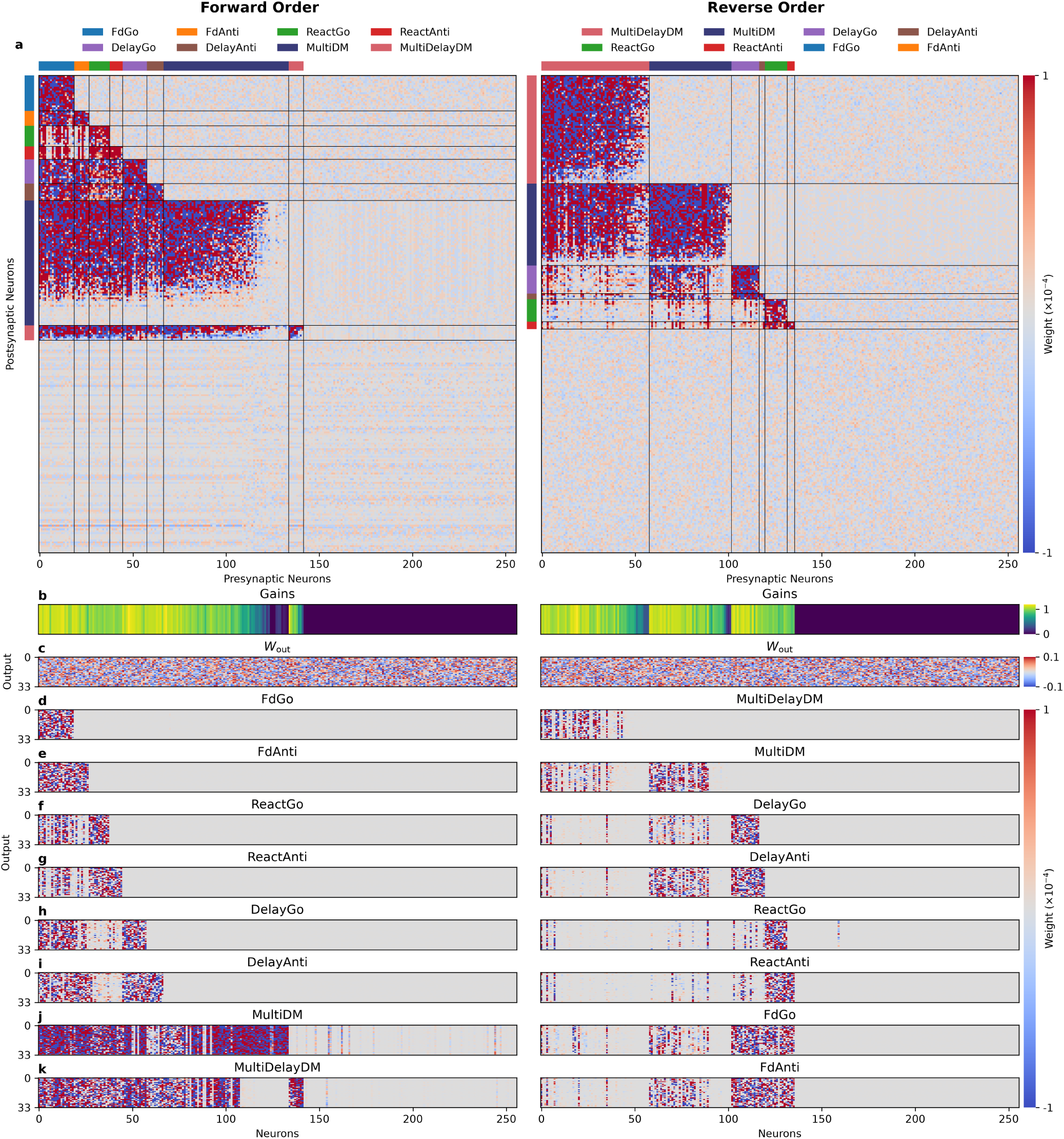
Same as Fig. 3, for random seed 128.

**Figure S4:**
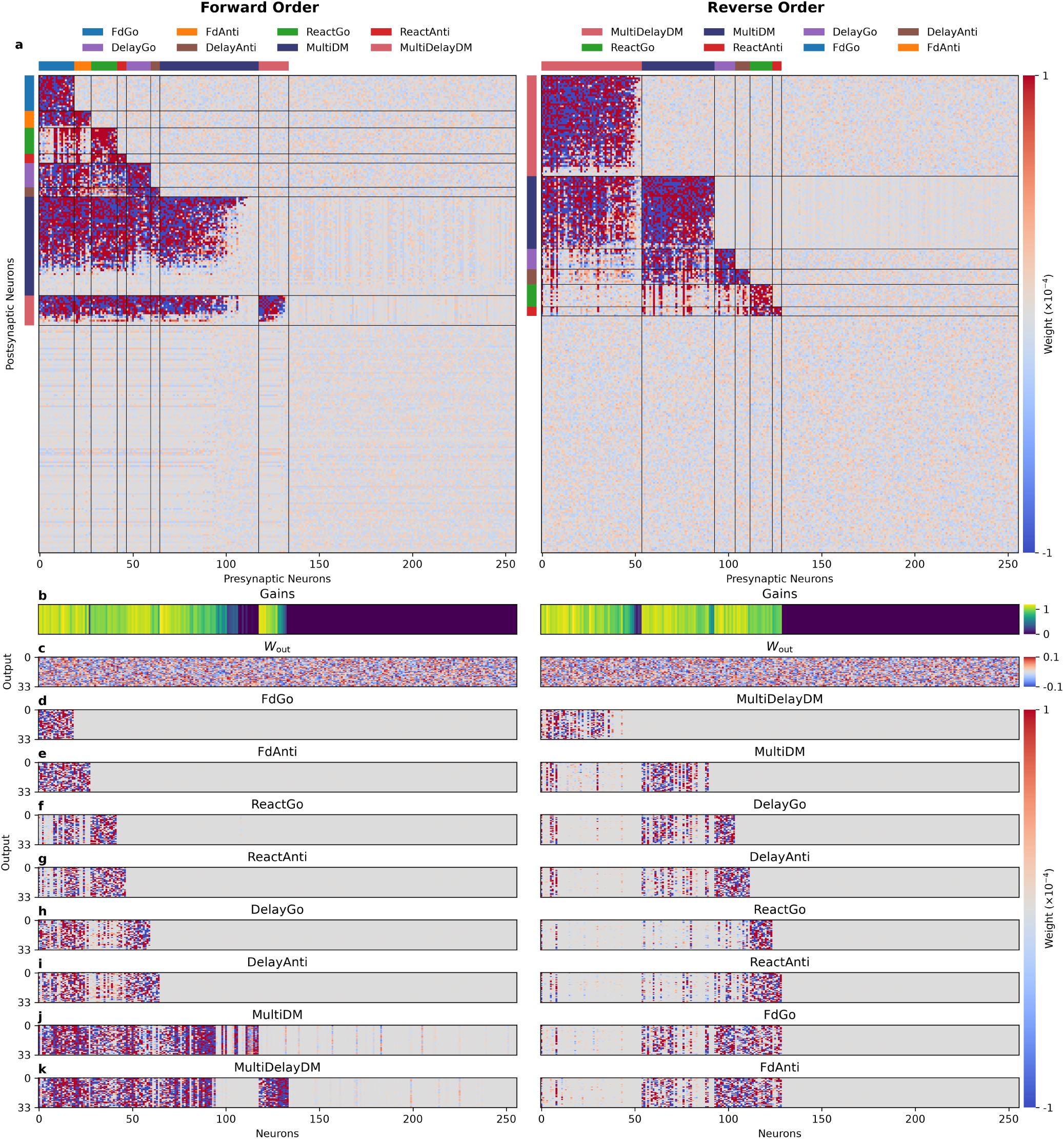
Same as Fig. 3, for random seed 128128.

**Figure S5:**
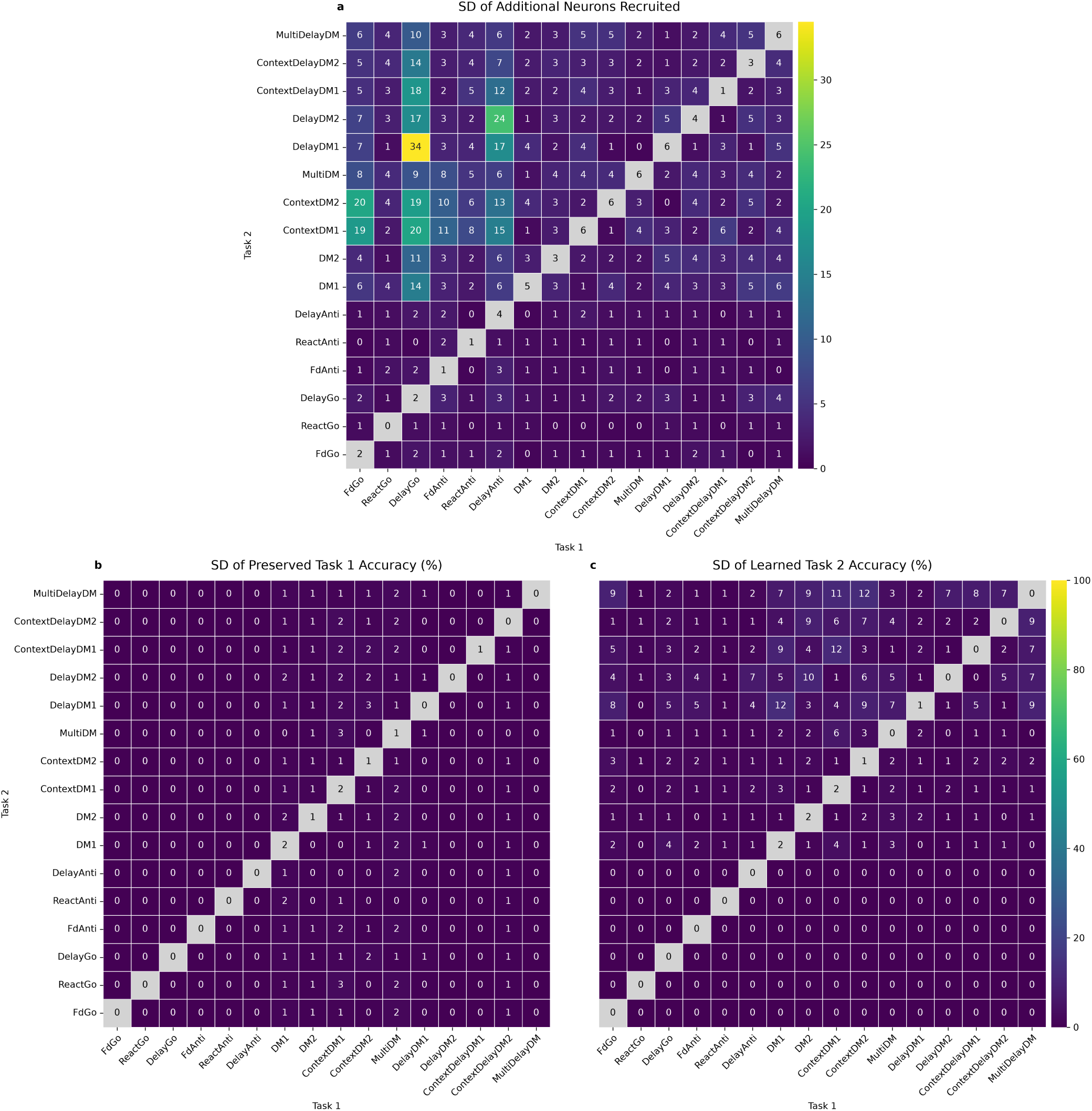
Pairwise recurrent allocation and task accuracy. Standard deviation (SD) across 4 random seeds for the means reported in Fig. 4. Columns denote the first learned task (Task 1), rows denote the second (Task 2). **a** SD of additional neurons recruited for Task 2. Diagonal entries are the SD of the number of neurons recruited by the task independently. **b** SD of preserved Task 1 accuracy after Task 2 training. Diagonal entries are the SD of accuracy learning task independently. **c** SD of Task 2 accuracy after its training. Diagonal entries are the SD of accuracy learning task independently.

**Figure S6:**
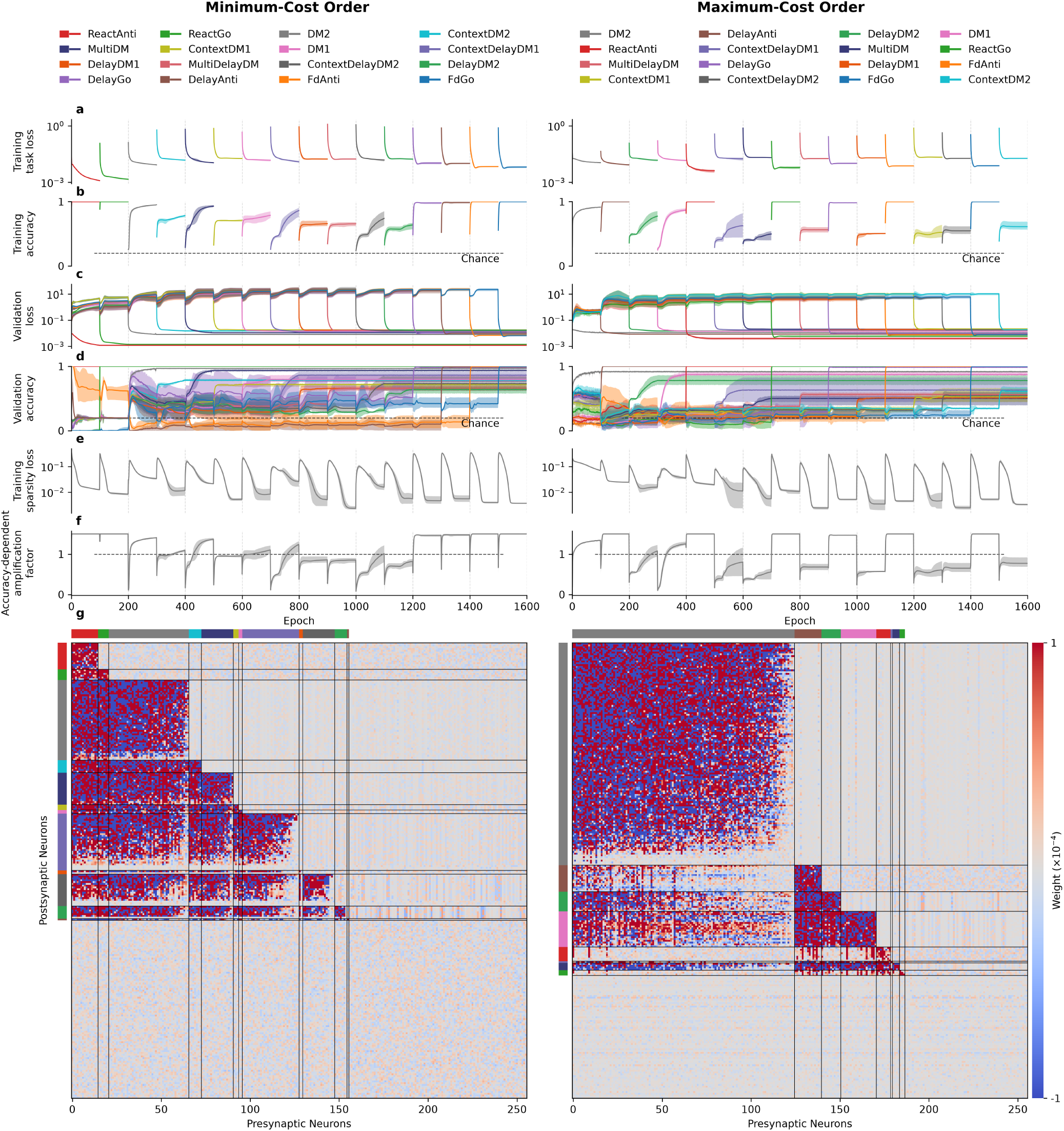
Same as Fig. 5, for random seed 42.

**Figure S7:**
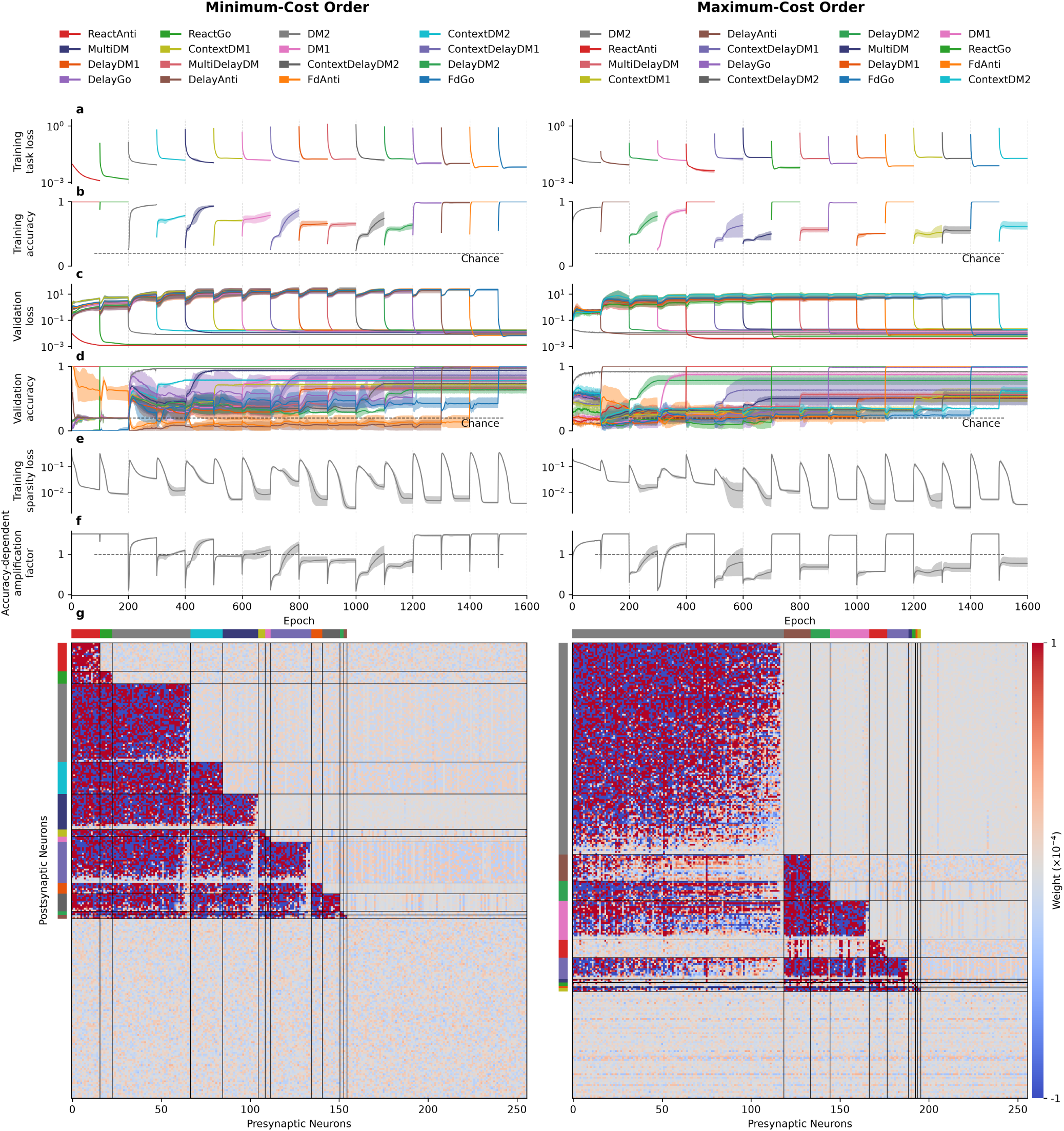
Same as Fig. 5, for random seed 128.

**Figure S8:**
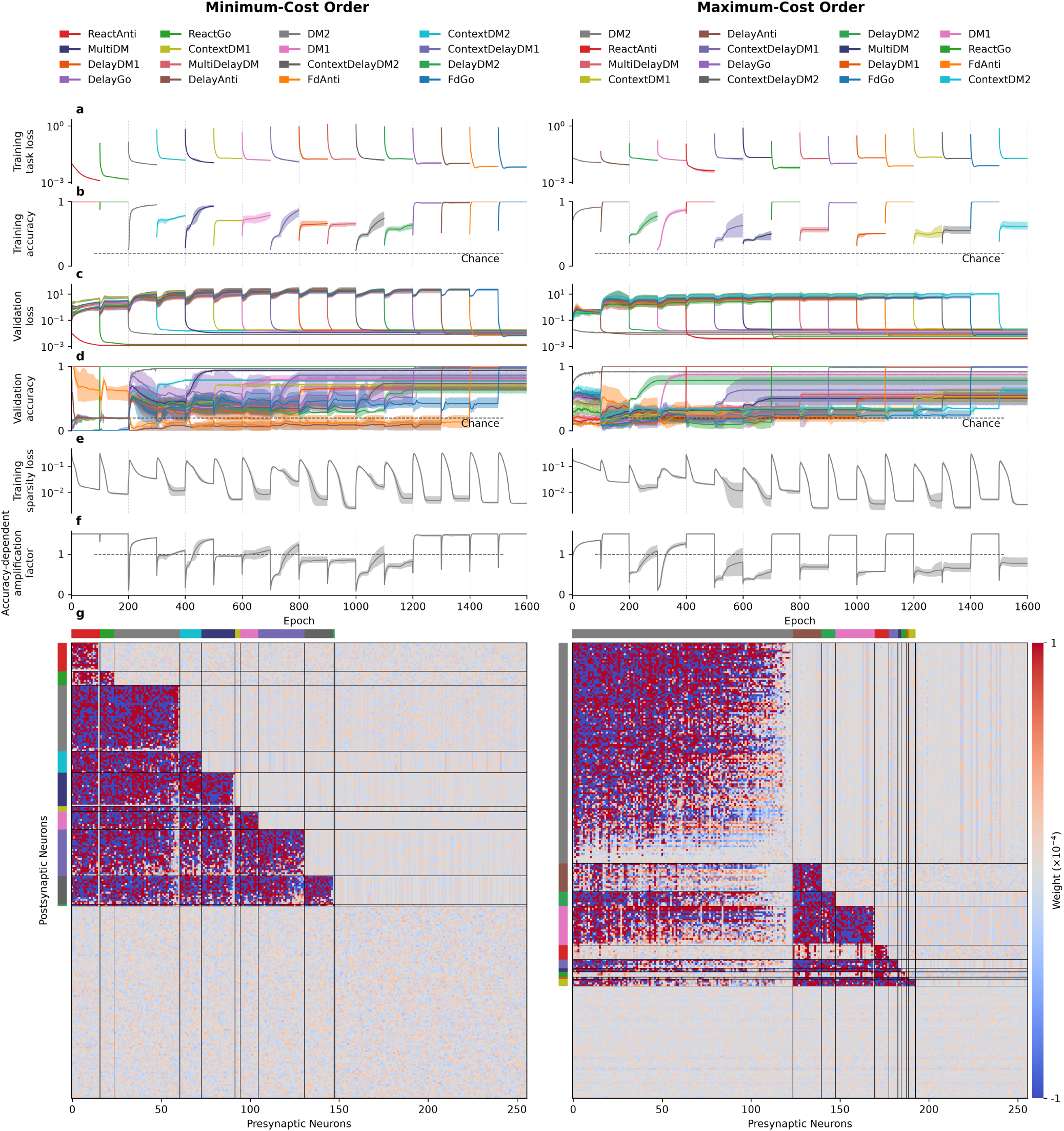
Same as Fig. 5, for random seed 128128.

